# Revisiting the landscape of evolutionary breakpoints across human genome using multi-way comparison

**DOI:** 10.1101/696245

**Authors:** Golrokh Vitae, Amine M. Remita, Abdoulaye Baniré Diallo

## Abstract

Genome rearrangement is one of the major forces driving the processes of the evolution and disease development. The chromosomal position affected by these rearrangements are called breakpoints. The breakpoints occurring during the evolution of species are known to be non randomly distributed. Detecting their landscape and mapping them to genomic features constitute an important features in both comparative and functional genomics. Several studies have attempted to provide such mapping based on pairwise comparison of genes as conservation anchors. With the availability of more accurate multi-way alignments, we design an approach to identify synteny blocks and evolutionary breakpoints based on UCSC 45-way conservation sequence alignments with 12 selected species. The multi-way designed approach with the mild flexibility of presence of selected species, helped to have a better determination of human lineage-specific evolutionary breakpoints. We identified 261,391 human lineage-specific evolutionary breakpoints across the genome and 2,564 dense regions enriched with biological processes involved in adaptive traits such as *response to DNA damage stimulus, cellular response to stress* and *metabolic process*. Moreover, we found 230 regions refractory to evolutionary breakpoints that carry genes associated with crucial developmental process such as *organ morphogenesis, skeletal system development, chordate embryonic development, nerve development* and *regulation of biological process*. This initial map of the human genome will help to gain better insight into several studies including developmental studies and cancer rearrangement processes.

## 1 Introduction

Genome rearrangement is one of the major forces driving the genome evolution, population diversity and development of diseases [10, 31, 35]. It happens when DNA breaks in specific positions (breakpoints) and reassembles in a way that results in a major transformation of genomic landscape [2]. It is now well-accepted that genome rearrangements are not random events and not all genomic regions have the same susceptibility to such drastic modifications [4, 19, 28, 29]. Millions of years of evolution and natural selection driven by these major structural modifications curved the genome in a way that regions carrying high functional and selective pressure maintained their integrity. Studies on the genome synteny have shown that conserved regions are not only significantly enriched in putative regulatory regions [18, 25] but also are associated with transcriptional regulations and developmental processes [25, 32, 37]. On the other hand, break-prone regions participate in speciation, adaptation and development of species-specific traits and behavior that are not detrimental to the survival nor reproduction of the species and known to carry distinctive functional and structural markers [6]. Although, the recent advancement in sequencing technology since the first draft of the human genome gave us an almost reliable version of the human genome in terms of its sequence, however, still less is known about the function of most genomic regions. Understanding the nature of the human genomic regions with respect to their susceptibility to genome rearrangements could give us an insight to the possible functionality of those regions and narrow down the search for further investigation. Hence, we based the main goal of this article to reveal the position of susceptible human genomic regions to large (over a certain size) rearrangements specific to human lineage. To identify the genomic regions subjected to rearrangements, first comes the identification of regions that have maintained their position, order and orientation (synteny blocks) as they should be conserved since a common ancestor and present in (relatively close) contemporary genomes. At that point, regions bordered by synteny blocks could be identified as genomic regions that have been subjected to rearrangements (breakpoint regions (BPRs) or breakpoints for short). However, the identification of synteny blocks are difficult and controversial due to several assumptions [16]. Selected species, type of comparison (pair-wise or multi-way), choice of conserved markers (anchors), minimum accepted gap between consecutive anchors and minimum anchor size, could all affect the position and size of synteny blocks therefore, the size and the position of the breakpoint regions [16, 33]. To identify human-lineage-specific synteny breaks, we developed a pipeline to extract synteny blocks based on the protocol of GRIMM-synteny [30]. However, to lower the bias caused by pair-wise comparison and genes as conservation anchor, we adjusted their method and applied it on a multi-way comparison of conserved genomic segments among a subset of selected species. Moreover, by using a human oriented multiple sequence alignments and having at least two species selected, per 4 major subset of mammalian phylogeny tree, we could eliminate the synteny breaks specific to non-human lineages within the mammalian reference tree. Furthermore, using a permutation test, we Identified the hotspots of breakpoints and identified their associations with 20 known structural and functional annotations.

## 2 Results

### 2.1 Identification of human lineage-specific breakpoint regions

In this study we aimed to identify the genomic regions maintained their synteny in the trajectory towards the evolution of the human genome along with their intermediary regions where rearrangements dynamically broke the synteny and drove the divergence and spoliation of the human species. Thus, to identify the synteny blocks, we applied our synteny block extraction method (4.2) to multiple conservation alignments blocks (multiz46way) of 45 vertebrates [12], given twelve selected species S3 and their phylogenetic tree. The procedure selects 21,809,215 blocks conserved among at least seven selected species. Each two neighboring anchors then were compared with respect to each species to identify the list of species with any synteny discontinuity (chromosome, orientation or distance*> δ*). In case of no discontinuity or discontinuity that have not been originated from any human ancestry nodes, the two neighboring anchors were fused together to form a cluster. The resultet 346,151 clusters then filtered by size (*<* 1*Kbp*) as the final step of synteny extraction. As a result 261,420 synteny blocks were identified on the human genome. These blocks cover 52% of the genome with a size distribution of 1 to 200 Kbp (*µ* : 6220.62, *σ* : 6958.4, *Md* : 3936). As explained before, a breakpoint is region between two successive synteny blocks. Therefore, all regions separating synteny blocks were considered as synteny breaks or breakpoint. It should be noted that these regions were passed through a filtering step and regions with size greater than 2 Mb were excluded from these regions. This constraint is necessary to avoid ambiguous regions that could not be assigned with high confidence to BPRs (e.g. sequences of heterochromatin). In the filtration step, the eliminated regions were mostly located on centromeric or telomeric regions and the rest were located on chromosome Y. The remaining 261,391 regions were identified as human lineage-specific breakpoints or BPRs. These BPRs cover 39.58% of the human genome. However, the coverage of these BPRs raise to over 40% in chr11, chr12, chr8, chr4 and chr7 and to over 55% in chr19 and chrX. See supplementary figure **??**. The size of these regions were from 1 bp to 1.857 Mbp. The seven largest BPRs (size *>* 1 Mbp) were located on chr14, chrY, chr11, chrX, chr2 and chr7. Based on a Least Common Ancestor approach, over 50% of these BPRs were reused and originated from separation of placental mammals. The next most representative ancestry node was the Boreotherian ancestor as the origin of 20% of BPRs. The distribution of BPRs ancestry origins is presented in the supplementary figure S2.

### 2.2 Localisation of breakpoint hotspots

Using an non-overlapping sliding window approach of size 100 Kbp and a permutation test (see 4.3), breakpoints were simulated based on their chromosome and size along the genome for 100,000 times. By comparing the breakpoint with simulated breakpoint counts in each window, a p-value were calculated. 2,564 window were identified as breakpoint hotspot regions (BHRs) having p-value *<* 0.05. Although evolutionary breakpoints are distributed across 90% of genomic windows, only 8.28% of genomic windows (2,564) were enriched in BPRs. These hotspots contain 12 to 21 breakpoints with an average of 14.85 *±* 1.56 (except for chromosome Y). BHRs located on chromosome Y harbor 3 to 14 breakpoints in each window. The BHRs distribution with respect to chromosomes shows that chromosomes are heterogeneously affected by these hotspots and the hotspot coverage varies from 3% to 23% among the chromosomes. The highest presence of hotspots were belong to chr19, chr15, chr21 and chr22 with a coverage over 18%.

### 2.3 The most resistant genomic regions to rearrangements

We found that less than one percent of genomic regions of 100 Kbp (230 regions) have a complete absence of breakpoints. It should be noted that more than 1400 windows with zero breakpoint have been eliminated as they were located on heterochromatin regions. These regions that are mostly continuous, located in 50 genomic locations on all chromosomes. We called these regions breakpoint deserts or breakpoint refractory regions (BRRs). The distribution of these regions are also varies among chromosomes. The 5 largest consecutive breakpoint deserts (from 1.7 Mbp to 2.8 Mbp) are located on chrY and chr19. See figure 1. It should be noted that more than half of these refractory regions (120 windows) do not overlap with any annotated genes. However, the remaining regions carry genes with high selection pressure conserved among primates, mammals or vertebrates. For instance, thirteen members of Hox-cluster genes that encode conserved transcription factors in most bilateral species [27] and 33 members of zinc finger genes located along almost complete chromosome band 19p12 (4.4 Mbp with one single breakpoint). These genes are known to be arisen in early primate evolution about 50 million years ago [1, 9]. The remaining genes presented in these breakpoint deserts were also conserved among mammals or vertebrates and any mutation in these genes could cause major developemental or reproduction effect on individuals. BCL11A, that mutation in this gene is known to be associated with intellectual developmental disorder with persistence of fetal hemoglobin (IDPFH) [26], SRY (sex-determining region Y), known to be associated with 46,XX testicular disorder of sex development and Swyer syndrome [26], FOXP1, known to be associated with Mental retardation with language impairment and autistic features (MRLIAF) [26], TCF4 known to be associated with Pitt-Hopkins syndrome is a condition characterized by intellectual disability and developmental delay, breathing problems, recurrent seizures (epilepsy), and distinctive facial features [26], are some examples of these genes and their importance in terms of survival and reproduction.

**Figure 1.**
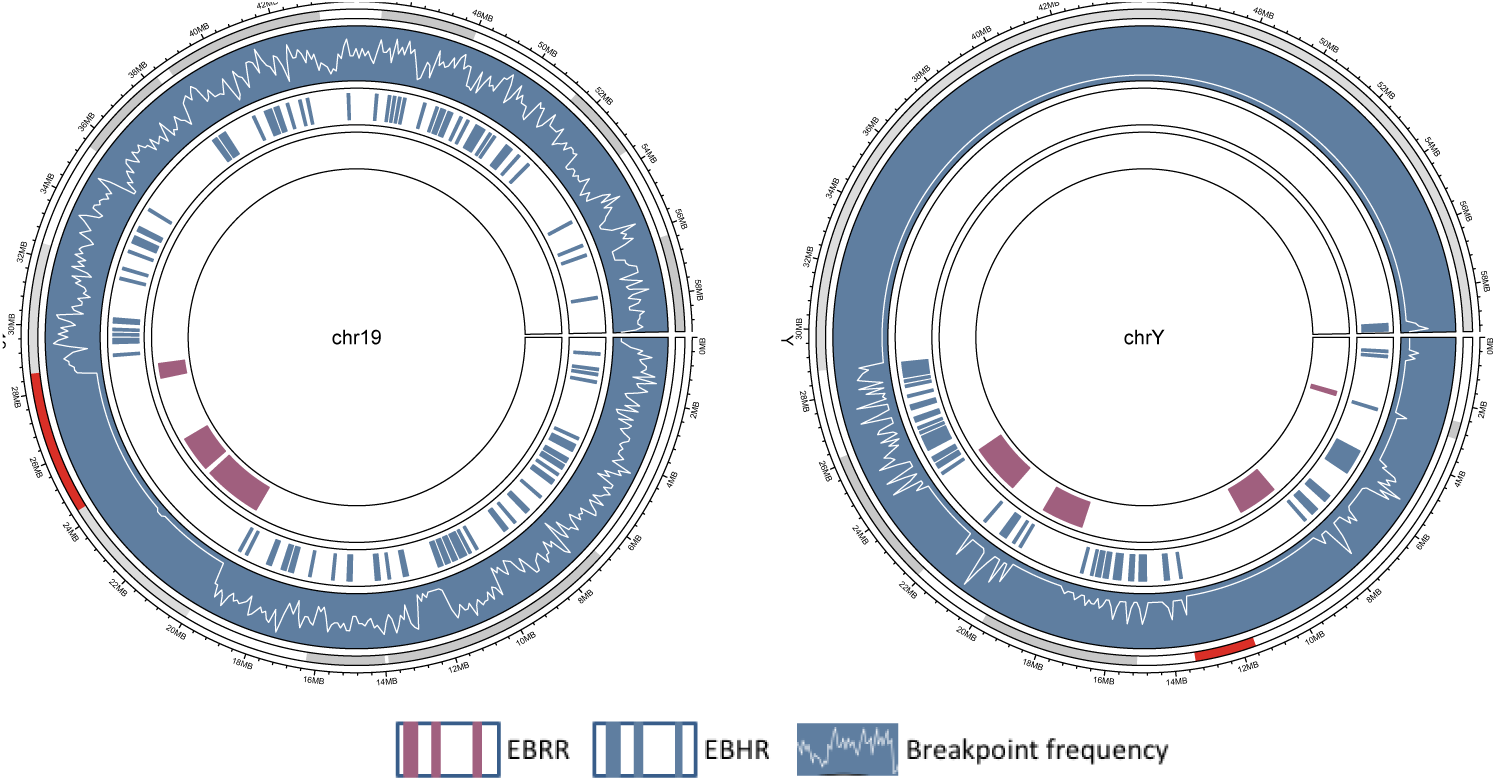
Chromosome 19 showed only one single BPRs in a complete chromosome band of 4.4 Mbp. This region codes for 33 members of zinc finger family which is highly conserved in primates. Chromosome Y is one of the most remarkable chromosomes on human genome. The smallest human chromosome with the highest presence of BPRs, BHRs and BRRs.

### 2.4 Functional enrichment of genomic regions

A GO enrichment analysis of biological process were performed to identify and compare the functional characteristics of different type of genomic regions with respect to their susceptibility to rearrangements. These results show that breakpoint deserts and breakpoint hotspot regions are enriched in distinct biological processes. Breakpoint hotspot regions are enriched in biological process associated with **response to stress** such as *response to DNA damage stimulus* (p-value = 9.1683e-4), *cellular response to stress* (p-value = 7.0041e-3), and *DNA repair* (p-value = 6.6835e-3); **metabolic process and transformation of chemical substances** such as *nucleic acid metabolic process* (p-value = 1.1152e-3), *macromolecule metabolic process* (p-value = 7.0041e-3) and *primary metabolic process* (p-value = 1.6258e-5); **chemical reactions and pathways** such as *catabolic process* (p-value = 5.9183e-3) as well as *cellular process* (p-value = 2.0792e-15). On the other hand, genes located within breakpoint deserts are enriched in biological processes completely different from breakpoint hotspots. These regions were highly enriched in critical biological processes **involved in development and morphogenesis** such as *anatomical structure arrangement* (p-value = 8.0323e-3), *anterior/posterior pattern formation* (p-value = 8.6894e-12), *chordate embryonic development* (p-value = 3.2238e-6), *cranial nerve morphogenesis* (p-value = 3.2706e-4), *embryonic development* (p-value = 1.0968e-4), *embryonic morphogenesis* (p-value = 9.8851e-5), *skeletal system morphogenesis* (p-value = 1.3727e-7), *nerve development* (p-value = 4.7057e-3), *organ morphogenesis* (p-value = 3.4475e-3) and **regulatory processes** such as *regulation of cellular metabolic process* (p-value = 2.3284e-16), *regulation of gene expression* (p-value = 3.5008e-20), *regulation of metabolic process* (p-value = 2.0673e-15), *regulation of transcription* (p-value = 3.9592e-22), etc. (See supplementary table S5 for the complete list of enriched biological process.)

### 2.5 Genomic features association

We studied the association of more than 20 genomic features with different identified genomic regions, BHRs, BRRs and others (the remaining). The coverage of each window for each genomic feature were calculated (BHR, BRR and other). Using a logistic regression we evaluated the importance of coverage of each genomic marker in prediction of the category of each genomic windows. An L2 regularization was used to ameliorate the prediction model. The results show that this logistic regression could classify the genomic regions based on the coverage of the selected features with an score of 0.90. These features have distinctive presence in each class of genomic regions (BRRs, BHRs and Others). These differences are illustrated in figure 2. Each cell corresponds to coefficient value calculated by the logistic regression for each class. For instance, conserved elements in Amniota (CEGA Amniota), Self chains and Low Complex Repeats have highest positive coefficient values for the class breakpoint desert (BRR) whereas for the class EBR Self chain has almost no effect (coefficient around zero) and high negative value for the class Other (non-BRR and non-BHR regions). G-quadruplex, genes, exons, benign structural variations, Common Fragile Sites, Synteny blocks, SINEs, Direct repeats and Mirror repeats ave high negative coefficient value for the class BRR where as positive coefficient for the other two classes. Another example is Simple and Tandem repeats with highest positive coefficient values for the class BHR but almost zero for the class BRR. The values and ranking of each genomic feature in this logistic regression classification are available in Supplementary Table S7

**Figure 2.**
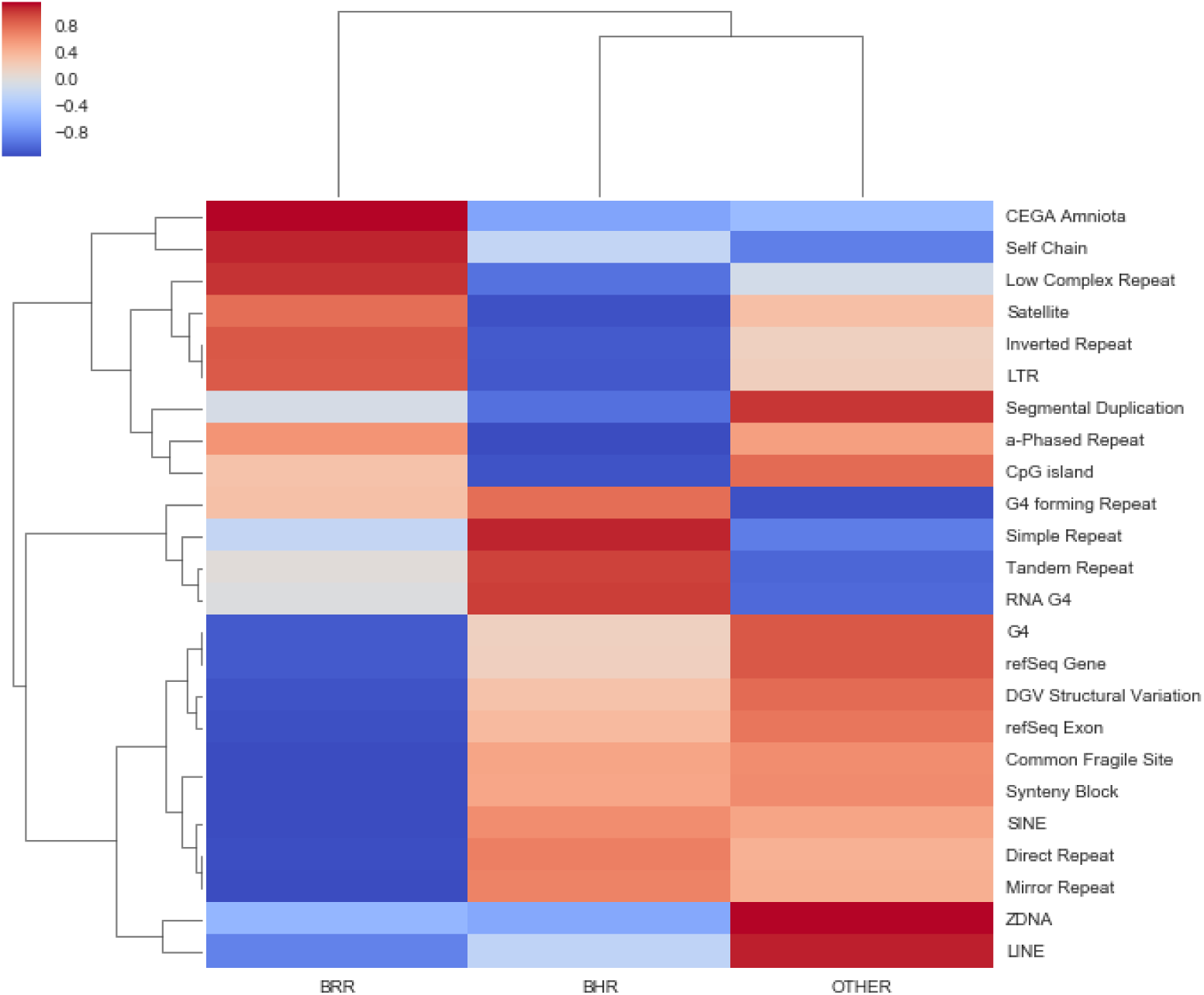
This figure shows the comparison between the coverage of selected genomic features on genome, BHRs and BRRs. x-axis represents the coverage percentage of each feature.

**Figure 3.**
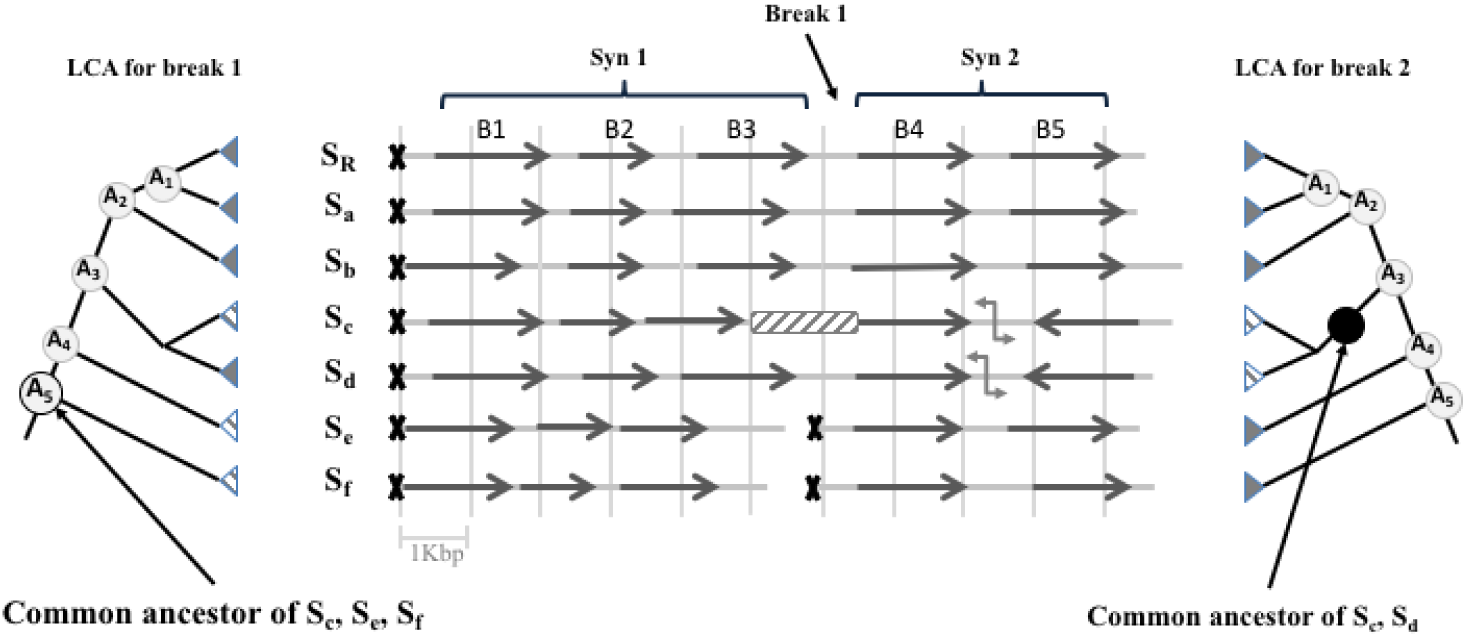
Extraction of synteny blocks and their corresponding breaks: In this illustration of a multi-way alignment blocks with respect to a reference species (S_*R*_). S_*a*_ to S_*f*_ are the selected species present in the alignment blocks. The reference tree show the phylogenetic relationship of these species. A_1_ to A_5_ are the S_*R*_ ancestry nodes. Each arrow represents a conserved region on each genome. Directions in each arrow represent the orientation of that region in each species. The distance between each two vertical lines represents 1 Kbp. Each X shows the start of a chromosome. The Break 1 is the result of the distance between B3 and B4 on S_*c*_ and change of chromosome in S_*e*_ and S_*f*_. The lowest common ancestor of these species is A_5_ which is an ancestor of S_*R*_. However, the second break in comparison between B4 and B5 caused by orientation change in S_*c*_ and S_*d*_ and their common ancestor is somewhere after separation of their clade. Hence, this break is not going to be counted as a break specific to the S_*R*_ lineage and B4 and B5 will be fused together.

## 3 Discussion

An evolutionary breakpoint region is a region that have been subjected to one or more rearrangement incident in the course of the evolution. Hence, to detect the position of such regions, first comes the identification of those regions that left intact since they are detectable as conserved regions among a group of relatively closed species (synteny blocks). Consequently, regions bounded by these conserved regions, are those that underwent a rearrangement incident also called breakpoint regions.

### 3.1 Evaluation of synteny identification

We conducted a multi-way comparative analysis of 12 genomes and identified synteny blocks and their corresponding 261,391 breakpoints (BPRs) on the human genome. Using a permutation approach, 2,564 regions of 100 Kbp were identified as breakpoint hotspot regions (BHRs). One of the basics of comparative genomics is the identification of synteny. However, there is no precise definition of synteny. Species selection, their evolutionary distances, pairwise or multi-way comparison, size of the anchors could all have huge impacts on the resulting synteny blocks (Reviewed by [16]). In this study 12 species were selected with the human as the reference species from different principal branches of species tree. We selected species according to their evolutionary distance from the human as well as the quality of their genome assembly. Species from more distant lineages of mammalian evolutionary tree and a non-mammalian vertebrate as out-group allows a higher resolution. However, it yields to shorter size of synteny blocks compared to previous studies. From multi-way alignment blocks, blocks shared by at least 7 selected species were accepted as conservation anchors. This looseness lowers the dependency of this method to the precise species list and could lower the bias of species selection and missing data. Also, the fact that about 80% of our BPRs have a Least Common Ancestor originated from the primate node to farther up on the reference tree, shows that these breakpoints are reuse and a careful replacement of some selected species should not have a dramatic effect on the identified synteny blocks and their positions. Nonetheless, comparing identified synteny blocks with Conserved Ancestral Regions (CARs) that reconstructed by [21], showed that 98.4% of identified synteny blocks overlaps with 59.98% CARs. Considering the estimation of [21] closest to the real ancestral genomic regions, the presence of synteny blocks in conserved ancestral regions is not far from the presence of these synteny regions in the human contemporary genome calculated in this study (52%). Due to the use of human oriented multi-way conservation alignments as anchors, over 7% of BPRs have a size equal to 1 bp as conservation anchors are contiguous on human genome. Comparison of BPRs with breakpoints produced in the study of [20] showed that BPRs were similar to what [20] identified as BPRs. It should be noted that, in this study pairwise comparison of orthologous genes based on five other mammals (with 4 that were common with species in this study) were used to identify synteny and their BPRs have similar size distribution of BPRs produced in our study. Other than the choice of the species, the use of genes as conservation anchors limited their study to only the coding regions. However, still 3% of BPRs produced in this study overlaps with 69.8% of their breakpoint regions.

### 3.2 Location of genes based on their different selective pressure

The results presented in this study highlights the dynamic nature of the genome. BPRs are dense in regions coding for functions related to adaptive response such as *cellular process, metabolic process, DNA repair, response to DNA damage stimulus, cellular response to stress* and *catabolic process*. On the other hand, genes with high selection pressure such as members of homeobox gene family of transcription factors, sex determining region Y (SRY), BAF chromatin remodeling complex subunit (BCL11A), Forkhead box transcription factors (FOXP1) and zinc finger gene family are located in EBRRs. All the genes in these regions are conserved among mammals and vertebrates. SRY initiates male sex determination. Mutation in this gene is known to be associated with 46,XY complete gonadal dysgenesis or 46,XY pure gonadal dysgenesis or Swyer syndrome [26]. Translocation of part of the Y chromosome containing this gene to the X chromosome causes XX male syndrome [15]. The homeobox genes encode a highly conserved family of transcription factors that play an important role in morphogenesis in all multicellular organisms [15]. HOXB genes are known to encode conserved transcription factors in most bilateral species [27]. FOXB1 belongs to subfamily P of the forkhead box (FOX) transcription factor family and known to be associated with Mental retardation with language impairment and autistic features (MRLIAF). Zinc finger genes code for transcription factors in all eukaryotes. 33 members of zinc finger genes located on the 19p12 chromosome band. This 4.4 Mbp region is the largest breakpoint desert in our dataset having only one single BPR within this region. This gene cluster is known to be highly conserved in primate [1, 9]. This region is presented in figure 1. These results are in harmony with previous study indicating that refractory regions are strongly enriched for genes involved in (embryonic) development [3, 11, 24] and any disruption in these regions should have detrimental effect [34]. Whereas, regions with high tendency of rearrangements are more involved in adaptive process such as metabolic process, DNA repair and response to stress [24].

In conclusion the identified BHRs provides a better insight on the nature of genome by emphasizing on the difference of the functions that breakpoint hotspots and breakpoint refractory regions harbor. It points out the dynamic process of evolution, adaptation and natural selection in interaction with the selective pressures of several genomic regions that maintain the integrity functions crucial for the survival of the species. The BHRs and BBRs reported in this paper will constitute a good asset for further studies including the affinity of some genomic regions to rearrangement associated to diseases known to be driven by genome rearrangements such as cancers.

## 4 Materials and Methods

### 4.1 Basic definitions

- ***Multi-way conservation alignment:*** is a series of multiple sequence alignments of genomic blocks on the human genome conserved among set of species. These alignment blocks are organized with respect to a single reference genome. The multiple sequence alignments used in this study were generated using multiz and other tools in the UCSC/Penn State Bioinformatics comparative genomics alignment pipeline. Conserved elements are identified by phastCons [17].
- ***Synetnic block:*** A syntenic block is defined as a large region of the genome that corresponds to a collection of contiguous blocks that maintained their positions and orders among a group of species since a common ancestor.
- ***Synteny break*** : Given two consecutive conserved synteny blocks, any discontinuity (as in distance, chromosome or orientation) between two sequences of one or more compared species that originated from any ancestry node of the reference genome (by Lowest Common Ancestor (LCA) approach) is considered a synteny breaks.
- ***Breakpoint region or simply breakpoint:*** genomic regions that is bounded by two neighboring syntenic blocks on the reference genome.

### 4.2 Synteny break identification

In order to identify the position of synteny breakpoints first comes the identification of synteny blocks as these regions maintained their location among a group of relatively close contemporary species since a common ancestor. To perform a comparative analysis on the human genome, the most appropriate candidate species to capture more evolutionary patterns and lower the bias produced by missing information, assembly errors and alignment errors in the analysis are of two kinds : 1) extremely close species that share common features, and 2) more or less distant species that could have a broad divergence with the reference species (human). Also, to enhance the prediction of breakpoint origin and be able to identify breakpoints that are not specific to human lineage, it is important to have at least two species present in each major clades of the reference species tree. Hence, to conduct our comparative analysis three well-studied primates have been chosen, chimpanzee, orangutan and marmoset for the first group and as for the second group, two well-sequenced and well-studied species from three major branches of mammalian phylogenetic tree were selected as follows: rat and mouse from Supraprimates, dog and cow from Laurasiatheria as well as elephant and armadillo as non-Boreotherian mammals. We also added two outliers, one non placental mammal, opossum and one non mammalian vertebrate, chicken.

Considering that we are looking for the conserved elements on the human genome to construct our synteny we decided to use multiple sequence alignments of conserved elements of 45 vertebrates with human as the reference species. The alignments are made available by UCSC genome browser in MAF format from http://hgdownload.cse.ucsc.edu/goldenPath/hg19/multiz46way/maf. Conservation alignment blocks having at least seven species among the 12 selected species were extracted as conservation anchors. Unselected species were removed from the blocks and missing selected species were added having unknown chromosome (chrUn) and zero as their positions. From there on synteny is constructed as follows.

Given:

- a set of selected species: *S* = *{s*_*R*_, *s*_*a*_, *s*_*b*_, *…, s*_*n*_*}*, with *s*_*R*_ as reference species.
- a species reference tree *T* with set of ancestry nodes of s_*R*_ *A* = *{A*_1_, *A*_2_, *A*_3_, *…, A*_*n*_*}*.
- Previously produced set of conservation anchors *B* = *{B*_1_, *B*_2_, *B*_3_, *…, B*_*n*_*}* each species sequence in a block *B*_*i*_ identified by its chromosome, *B*_*i*_.*s*_*c*_, its orientation, *B*_*i*_.*s*_*o*_ and its head *Bi.s*_*h*_ and tail *B*_*i*_.*s*_*t*_

*\*Note:* Discontinuity between two blocks, *B*_*i*_ and *B*_*j*_ with respect to each species is defined based on any of the following three conditions:

- difference in chromosomes: *B*_*i*_.*s*_*c*_ *≠ B*_*j*_.*s*_*c*_
- difference in orientations: *B*_*i*_.*s*_*o*_ *≠ B*_*j*_.*s*_*o*_
- distance, *δ*(*B*_*j*_.*s*_*t*_ *− B*_*i*_.*s*_*h*_), greater than a defined threshold *G*: *δ > G*

**For each** two consecutive conservation anchors, *B*_*i*_ and *B*_*j*_:

1. S^*\**^ *←−* get subset of *S* with discontinuity (*B*_*i*_, *B*_*j*_)
2. if LCA(S^*\**^) *⊂ A {Synteny is broken along the evolution of reference species genome}*then:

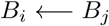
3. else

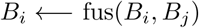
4. *B*_*j*_ *←− B*_*j*+1_.

For each two continuous anchors (with respect to the reference genome), species with any discontinuity (chromosome, orientation and distance) were identified. If the lowest common ancestor of species with discontinuity is an element of human ancestry nodes of the reference species tree, the two blocks would be considered as discontinuous and the algorithm will continue to compare the next couple in the line. Otherwise, the two blocks will be fused together to be compared with the next block. In each iteration, the list of species with discontinuity and their LCA will be documented. The results of this step is a list of larger conservation clusters that will go through a size filter (minimum size of 1000 bp). The clusters passed through the filter are the resulting synteny blocks. These steps are well illustrated in figure **??**. As previously noted, a breakpoint region is a region bounded by two consecutive synteny blocks. However, these regions passed through a size filtering step and only regions with a size smaller than 2 Mbp are considered as reference species lineage-specific breakpoints, synteny breaks, evolutionary breakpoints or breakpoint regions (BPRs). The size constraint is necessary to avoid ambiguous regions that could not be associated correctly to breakpoints (e.g. sequences of heterochromatin). The origin of each BPR is the lowest common ancestor of species with discontinuity comparing the two bounded synteny blocks in the synteny identification step.

### 4.3 Prediction of breakpoint-hotspots

To identify the genomic regions that are significantly enriched in breakpoint or breakpoint hotspot regions, we followed the permutation strategies suggested by *De et al.* 2013[7]. Genome was scanned with a non-overlapping sliding window of size 100 Kb and the number of breakpoints for each window was counted. Breakpoints were considered to fall into a window if they have at least one position overlapping that window. For 100,000 iterations, each BPR in is simulated based on the following constraints:

- Breakpoint should be simulated within the original chromosome.
- Simulated breakpoint should not fall into centromeric or telomeric regions.
- Breakpoint should be simulated with respect to its size.
- Overlaps would be allowed to lower the calculation costs.

A p-value is calculated for the number of simulated breakpoints per window with compare to the original count. Windows with a p-vaule ¡ 0.05 is considered as significantly enriched windows or breakpoint hotspot regions (BHRs).

### 4.4 Collection of genomic features

CpG islands, nested repeats, segmental duplications, CG content and self chain markers were downloaded from the UCSC Genome Browser [23]. Annotation on non-B DNA conformation were downloaded from Non-B DB [5]. Conserved elements of amniotes were obtained from catalog of conserved elements from genomic alignments (CEGA) [8]. Common fragile sites were provided by the supplementary material of [13] paper. RNA G-quadrupex were downloaded from RNA G-quadruplex database (G4RNA) [14]. Benign structural variation were donwloaded from Database of Genomic Variants [22]. RefSeq genes and exons where obtained from NCBI FTP portal. G quadruplex were downloaded from G4 database [36] in 2013. However, the database is not accessible anymore. The complete list of these features are represented in supplementary table S6.

### 4.5 Identification of breakpoint deserts

Regions with complete absent of BPR were identified and their positions were compared to genomic regions that are not well sequenced or annotated. We found 1,462 genomic windows outside telomeric and centromeric regions that fall into other heterochromatin gaps such as short arms of chromosome 13, 14, 15, 21, 22, as well as chromosome bounds 1q12, 9q12 and 16q11. These regions were eliminated from our analysis. Hence 230 windows of 100 Kbp remained with no EBr. These regions are labeled as breakpoint refractory regions (BRRs) or break point deserts.

## Supporting Information

Supporting data for this study is available at:

https://github.com/bioinfoUQAM/RECOMB-CG-2019_supp

## Acknowledgments

We would like to thank Julie Horvath, Aida Ouangraoua, Abou Abdallah Malick Diouara and Bruno Daigle for helpful discussions. Special thanks to Emmanuel Mongin, by whom this project has been initially inspired. This work is supported by Natural Science and Engineering council of Canada (NSERC) and the Fonds de Recherche du Québec-Nature et Technologie (FRQNT) funds to ABD. AMR is a NSERC fellow. AMR and GV are FRQNT fellows….

## References

1. E. J. Bellefroid, J.-C. Marine, A. G. Matera, C. Bourguignon, T. Desai, K. C. Healy, P. Bray-Ward, J. A. Martial, J. N. Ihle, and D. C. Ward. Emergence of the znf91 krüppel-associated box-containing zinc finger gene family in the last common ancestor of anthropoidea. Proceedings of the National Academy of Sciences, 92(23):10757–10761, 1995.

2. M. Blanchette. Evolutionary puzzles: An introduction to genome rearrangement. In Computational Science-ICCS 2001, pages 1003–1011. Springer, 2001.

3. D. Boffelli, M. A. Nobrega, and E. M. Rubin. Comparative genomics at the vertebrate extremes. Nature Reviews Genetics, 5(6):456, 2004.

4. G. Bourque, P. A. Pevzner, and G. Tesler. Reconstructing the genomic architecture of ancestral mammals: lessons from human, mouse, and rat genomes. Genome research, 14(4):507–516, 2004.

5. R. Z. Cer, K. H. Bruce, U. S. Mudunuri, M. Yi, N. Volfovsky, B. T. Luke, A. Bacolla, J. R. Collins, and R. M. Stephens. Non-b db: a database of predicted non-b dna-forming motifs in mammalian genomes. Nucleic acids research, 39(Suppl 1):D383–D391, 2010.

6. S. De and F. Michor. Dna secondary structures and epigenetic determinants of cancer genome evolution. Nature structural & molecular biology, 18(8):950–955, 2011.

7. S. De, B. S. Pedersen, and K. Kechris. The dilemma of choosing the ideal permutation strategy while estimating statistical significance of genome-wide enrichment. Briefings in bioinformatics, 15(6):919–928, 2013.

8. A. Dousse, T. Junier, and E. M. Zdobnov. Cega—a catalog of conserved elements from genomic alignments. Nucleic acids research, 44(D1):D96–D100, 2015.

9. E. E. Eichler, S. M. Hoffman, A. A. Adamson, L. A. Gordon, P. McCready, J. E. Lamerdin, and H. W. Mohrenweiser. Complex *β*-satellite repeat structures and the expansion of the zinc finger gene cluster in 19p12. Genome research, 8(8):791–808, 1998.

10. E. E. Eichler and D. Sankoff. Structural dynamics of eukaryotic chromosome evolution. Science, 301(5634):793–797, 2003.

11. P. G. Engström, S. J. H. Sui, Ø. Drivenes, T. S. Becker, and B. Lenhard. Genomic regulatory blocks underlie extensive microsynteny conservation in insects. Genome research, 17(12):1898–1908, 2007.

12. P. A. Fujita, B. Rhead, A. S. Zweig, A. S. Hinrichs, D. Karolchik, M. S. Cline, M. Goldman, G. P. Barber, H. Clawson, A. Coelho, et al. The ucsc genome browser database: update 2011. Nucleic acids research, page gkq963, 2010.

13. A. Fungtammasan, E. Walsh, F. Chiaromonte, K. A. Eckert, and K. D. Makova. A genome-wide analysis of common fragile sites: what features determine chromosomal instability in the human genome? Genome research, 22(6):993–1005, 2012.

14. J.-M. Garant, M. J. Luce, M. S. Scott, and J.-P. Perreault. G4rna: an rna g-quadruplex database. Database, 2015, 2015.

15. L. Y. Geer, A. Marchler-Bauer, R. C. Geer, L. Han, J. He, S. He, C. Liu, W. Shi, and S. H. Bryant. The ncbi biosystems database. Nucleic acids research, 38(Suppl 1):D492–D496, 2009.

16. C. G. Ghiurcuta and B. M. Moret. Evaluating synteny for improved comparative studies. Bioinformatics, 30(12):i9–i18, 2014.

17. W. J. Kent, C. W. Sugnet, T. S. Furey, K. M. Roskin, T. H. Pringle, A. M. Zahler, and D. Haussler. The human genome browser at ucsc. Genome research, 12(6):996–1006, 2002.

18. H. Kikuta, M. Laplante, P. Navratilova, A. Z. Komisarczuk, P. G. Engström, D. Fredman, A. Akalin, M. Caccamo, I. Sealy, K. Howe, et al. Genomic regulatory blocks encompass multiple neighboring genes and maintain conserved synteny in vertebrates. Genome research, 17(5):545–555, 2007.

19. C. Lemaitre, L. Zaghloul, M.-F. Sagot, C. Gautier, A. Arneodo, E. Tannier, and B. Audit. Analysis of fine-scale mammalian evolutionary breakpoints provides new insight into their relation to genome organisation. BMC genomics, 10(1):335, 2009.

20. C. Lemaitre, L. Zaghloul, M.-F. Sagot, C. Gautier, A. Arneodo, E. Tannier, and B. Audit. Analysis of fine-scale mammalian evolutionary breakpoints provides new insight into their relation to genome organisation. BMC genomics, 10(1):335, 2009.

21. J. Ma, L. Zhang, B. B. Suh, B. J. Raney, R. C. Burhans, W. J. Kent, M. Blanchette, D. Haussler, and W. Miller. Reconstructing contiguous regions of an ancestral genome. Genome research, 16(12):1557–1565, 2006.

22. J. R. MacDonald, R. Ziman, R. K. Yuen, L. Feuk, and S. W. Scherer. The database of genomic variants: a curated collection of structural variation in the human genome. Nucleic acids research, 42(D1):D986–D992, 2013.

23. L. R. Meyer, A. S. Zweig, A. S. Hinrichs, D. Karolchik, R. M. Kuhn, M. Wong, C. A. Sloan, K. R. Rosenbloom, G. Roe, B. Rhead, et al. The ucsc genome browser database: extensions and updates 2013. Nucleic acids research, 41(D1):D64–D69, 2012.

24. E. Mongin, K. Dewar, and M. Blanchette. Long-range regulation is a major driving force in maintaining genome integrity. BMC evolutionary biology, 9(1):203, 2009.

25. E. Mongin, K. Dewar, and M. Blanchette. Mapping association between long-range cis-regulatory regions and their target genes using synteny. Journal of Computational Biology, 18(9):1115–1130, 2011.

26. H. page: National Library of Medicine (US). Genetics Home Reference [Internet]. Bethesda (MD). The library. Internet, June 2019.

27. J. C. Pearson, D. Lemons, and W. McGinnis. Modulating hox gene functions during animal body patterning. Nature Reviews Genetics, 6(12):893, 2005.

28. Q. Peng, P. A. Pevzner, and G. Tesler. The fragile breakage versus random breakage models of chromosome evolution. PLoS computational biology, 2(2):e14, 2006.

29. P. Pevzner and G. Tesler. Genome rearrangements in mammalian evolution: lessons from human and mouse genomes. Genome Research, 13(1):37–45, 2003.

30. P. Pevzner and G. Tesler. Genome rearrangements in mammalian evolution: lessons from human and mouse genomes. Genome Research, 13(1):37–45, 2003.

31. L. H. Rieseberg. Chromosomal rearrangements and speciation. Trends in ecology & evolution, 16(7):351–358, 2001.

32. A. Sandelin, P. Bailey, S. Bruce, P. G. Engström, J. M. Klos, W. W. Wasserman, J. Ericson, and B. Lenhard. Arrays of ultraconserved non-coding regions span the loci of key developmental genes in vertebrate genomes. BMC genomics, 5(1):99, 2004.

33. D. Sankoff. The where and wherefore of evolutionary breakpoints. J Biol, 8:66, 2009.

34. F. Spitz, C. Herkenne, M. A. Morris, and D. Duboule. Inversion-induced disruption of the hoxd cluster leads to the partition of regulatory landscapes. Nature genetics, 37(8):889, 2005.

35. P. Stankiewicz and J. R. Lupski. Genome architecture, rearrangements and genomic disorders. TRENDS in Genetics, 18(2):74–82, 2002.

36. H. M. Wong, O. Stegle, S. Rodgers, and J. L. Huppert. A toolbox for predicting g-quadruplex formation and stability. Journal of nucleic acids, 2010, 2010.

37. A. Woolfe, M. Goodson, D. K. Goode, P. Snell, G. K. McEwen, T. Vavouri, S. F. Smith, P. North, H. Callaway, K. Kelly, et al. Highly conserved non-coding sequences are associated with vertebrate development. PLoS biology, 3(1):e7, 2004.

