## Supplementary for "Revisiting the landscape of evolutionary breakpoints across human genome using multi-way comparison"

### 5 Supplementary results

| Method | Input | Output | Size of output |
|---|---|---|---|
| Conservation anchor extraction | 45-way multiple alignments | Conservation anchors | 21,809,215 |
| Fusion | Conservation anchors | Synteny clusters | 346,515 |
| filter according to length | Synteny clusters | Synteny blocks | 261,420 |
| Synteny break extraction | Synteny blocks | Synteny breaks | 261,421 |
| Size filter | Synteny breaks | EBrs | 261,391 |
| Permutation | EBrs | EBHRs | 2,564 |
| EBr desert extraction | EBrs + human genome | EBRRs | 230 |

**Table S1.** Summary of EBr-related results

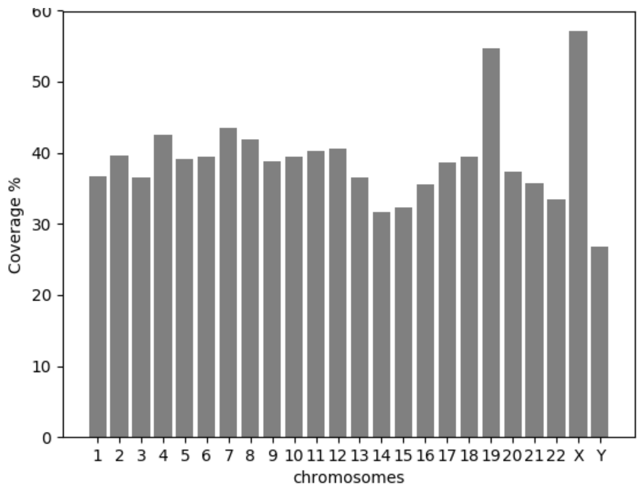


**Figure S1.** Coverage of chromosomes by EBrs

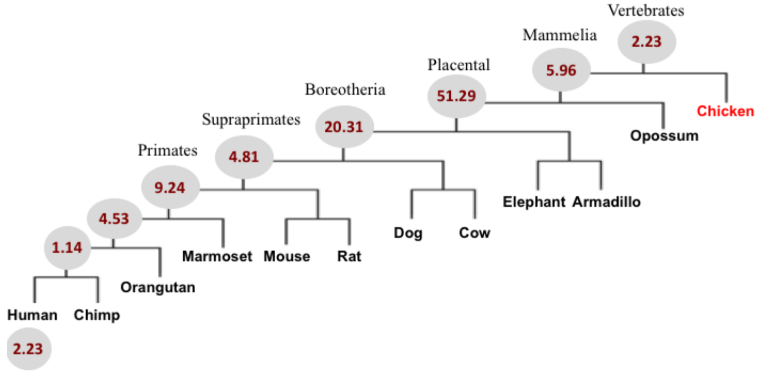


**Figure S2.** The percentages of human lineage specific breakpoints origin along the vertebrates reference tree

| GO id | Biological Process | adjt p-val | nb gene |
|---|---|---|---|
| GO:9987 | cellular process | 2.0792e-15 | 1464 |
| GO:44237 | cellular metabolic process | 4.5709e-7 | 814 |
| GO:8152 | metabolic process | 0.0000013601 | 946 |
| GO:44238 | primary metabolic process | 0.000016258 | 842 |
| GO:6807 | nitrogen compound metabolic process | 0.00071016 | 377 |
| GO:6974 | response to DNA damage stimulus | 0.00091683 | 89 |
| GO:44260 | cellular macromolecule metabolic process | 0.00091683 | 570 |
| GO:34641 | cellular nitrogen compound metabolic process | 0.00091683 | 357 |
| GO:6139 | nucleobase, nucleoside, nucleotide and nucleic acid metabolic process | 0.0011152 | 312 |
| GO:9056 | catabolic process | 0.0059183 | 186 |
| GO:6281 | DNA repair | 0.0066835 | 86 |
| GO:33554 | cellular response to stress | 0.0070041 | 68 |
| GO:43170 | macromolecule metabolic process | 0.0070041 | 633 |

**Table S2.** Biological process of Gene ontology enriched terms inEBHRs

| Species | Scientific Name | Assembly | nb nucleotide | alignment coverage | coverage of hg19 anchors |
|---|---|---|---|---|---|
| Human | Homo sapiens | hg19/GRCh37 | 1473659829 | 100 | 47.60 |
| Chimpanzee | Pan troglodytes | panTro2/Pan_troglodytes-2.1 | 1452392255 | 98.56 | 46.92 |
| Orangutan | Pongo abelii | ponAbe2/P_pygmaeus_2.0.2 | 1430816053 | 97.09 | 46.22 |
| Marmoset | Callithrix jacchus | calJac1/Callithrix jacchus-3.2 | 1376766952 | 93.42 | 44.47 |
| Mouse | Mus musculus | mm9/MGSCv37 | 1003857066 | 68.12 | 32.43 |
| Rat | Rattus norvegicus | rn4/RGSC_v3.4 | 938532785 | 63.69 | 30.32 |
| Dog | Canis lupus familiaris | canFam2/CanFam2.0 | 1320175342 | 89.58 | 42.65 |
| Cow | Bos taurus | bosTau4/Btau_4.0 | 1224047143 | 83.06 | 39.54 |
| Elephant | Loxodonta africana | loxAfr3/Loxafr3.0 | 1250650841 | 84.87 | 40.40 |
| Armadillo | Dasypus novemcinctus | dasNav2/Dasnov2.0 | 900944364 | 61.14 | 29.10 |
| Opossum | Monodelphis domestica | monDom5/MonDom5 | 342113074 | 23.22 | 11.05 |
| Chicken | Gallus gallus | galGal3/Gallus_gallus-2.1 | 86484085 | 5.87 | 2.79 |

**Table S3.** List of selected species and their contribution in extracted conserved anchors and the contribution of each genome in initial multiple sequence alignment of 45 species against human in nucleotide. Alignment coverage column carries information on the presence of a genome in the extracted conserved alignment blocks. The last column shows the contribution of each genome in the initial MSA in terms of nucleotide nucleotide.

| **chr** | **nbWind per chr** | **nb BHR** | **chr proportion** |
|---|---|---|---|
| chr4 | 1912 | 58 | 3.03 |
| chr3 | 1981 | 73 | 3.69 |
| chr6 | 1712 | 71 | 4.15 |
| chr20 | 631 | 28 | 4.44 |
| chr2 | 2432 | 121 | 4.98 |
| chr18 | 781 | 39 | 4.99 |
| chr5 | 1810 | 101 | 5.58 |
| chr10 | 1356 | 79 | 5.83 |
| chr7 | 1592 | 96 | 6.03 |
| chr8 | 1464 | 90 | 6.15 |
| chr12 | 1339 | 86 | 6.42 |
| chr17 | 812 | 59 | 7.27 |
| chr11 | 1351 | 108 | 7.99 |
| chr1 | 2493 | 204 | 8.18 |
| chr16 | 904 | 94 | 10.40 |
| chr13 | 1152 | 133 | 11.55 |
| chrY | 594 | 72 | 12.12 |
| chrX | 1553 | 193 | 12.43 |
| chr9 | 1413 | 183 | 12.95 |
| chr14 | 1074 | 156 | 14.53 |
| chr19 | 592 | 108 | 18.24 |
| chr15 | 1026 | 195 | 19.01 |
| chr21 | 482 | 97 | 20.12 |
| chr22 | 514 | 120 | 23.35 |

**Table S4.** BHR windows count across chromosomes.

| GO id | Biological Process | adjt p-val | nb gene |
|---|---|---|---|
| GO:6355 | regulation of transcription, DNA-dependent | 1.7061e-23 | 47 |
| GO:51252 | regulation of RNA metabolic process | 2.7934e-23 | 47 |
| GO:45449 | regulation of transcription | 3.9592e-22 | 52 |
| GO:19219 | regulation of nucleobase, nucleoside, nucleotide and nucleic acid metabolic process | 2.1082e-20 | 53 |
| GO:10556 | regulation of macromolecule biosynthetic process | 2.1082e-20 | 52 |
| GO:51171 | regulation of nitrogen compound metabolic process | 2.2033e-20 | 53 |
| GO:10468 | regulation of gene expression | 3.5008e-20 | 52 |
| GO:31326 | regulation of cellular biosynthetic process | 1.3681e-19 | 52 |
| GO:9889 | regulation of biosynthetic process | 1.7743e-19 | 52 |
| GO:60255 | regulation of macromolecule metabolic process | 2.1476e-17 | 52 |
| GO:80090 | regulation of primary metabolic process | 2.341e-17 | 53 |
| GO:31323 | regulation of cellular metabolic process | 2.3284e-16 | 53 |
| GO:19222 | regulation of metabolic process | 2.0673e-15 | 53 |
| GO:48706 | embryonic skeletal system development | 7.0326e-12 | 11 |
| GO:9952 | anterior/posterior pattern formation | 8.6894e-12 | 13 |
| GO:3002 | regionalization | 8.3379e-10 | 13 |
| GO:50794 | regulation of cellular process | 1.3375e-8 | 56 |
| GO:48704 | embryonic skeletal system morphogenesis | 1.3783e-8 | 8 |
| GO:7389 | pattern specification process | 1.5795e-8 | 13 |
| GO:48562 | embryonic organ morphogenesis | 3.8235e-8 | 10 |
| GO:50789 | regulation of biological process | 1.1284e-7 | 56 |
| GO:48705 | skeletal system morphogenesis | 1.3727e-7 | 9 |
| GO:1501 | skeletal system development | 1.5339e-7 | 13 |
| GO:65007 | biological regulation | 0.0000012646 | 56 |
| GO:48568 | embryonic organ development | 0.0000018797 | 10 |
| GO:43009 | chordate embryonic development | 0.0000032238 | 12 |
| GO:9792 | embryonic development ending in birth or egg hatching | 0.0000034982 | 12 |
| GO:48598 | embryonic morphogenesis | 0.000098851 | 10 |
| GO:9790 | embryonic development | 0.00010968 | 13 |
| GO:21602 | cranial nerve morphogenesis | 0.00032706 | 3 |
| GO:21570 | rhombomere 4 development | 0.001241 | 2 |
| GO:21545 | cranial nerve development | 0.0022559 | 3 |
| GO:21610 | facial nerve morphogenesis | 0.0022559 | 2 |
| GO:21612 | facial nerve structural organization | 0.0022559 | 2 |
| GO:21561 | facial nerve development | 0.003444 | 2 |
| GO:21604 | cranial nerve structural organization | 0.003444 | 2 |
| GO:60216 | definitive hemopoiesis | 0.003444 | 2 |
| GO:9887 | organ morphogenesis | 0.0034475 | 11 |
| GO:21675 | nerve development | 0.0047057 | 3 |
| GO:21546 | rhombomere development | 0.0063311 | 2 |
| GO:21783 | preganglionic parasympathetic nervous system development | 0.0063311 | 2 |
| GO:2011 | morphogenesis of an epithelial sheet | 0.0063311 | 2 |
| GO:48532 | anatomical structure arrangement | 0.0080323 | 2 |
| GO:48857 | neural nucleus development | 0.0080323 | 2 |

**Table S5.** Gene ontology enrichment analysis: Biological process enriched in EBRRs

| marker category | marker | source |
| --- | --- | --- |
| DNA conformation | a-phased_repeats | non-B database |
| | direct repeats | non-B database |
| | g-quadruplex_forming_repeats | non-B database |
| | inverted repeats | non-B database |
| | mirror repeats | non-B database |
| | short tandem repeats | non-B database |
| | Z-DNA | non-B database |
| DNA sequnece | LINE | UCSC nested Repeats |
| | Low complexity repeats | UCSC nested Repeats |
| | LTR | UCSC nested Repeats |
| | Satellite repeats | UCSC nested Repeats |
| | Simple repeat | UCSC nested Repeats |
| | SINE | UCSC nested Repeats |
| | SD | UCSC |
| | G quadruplex | G4 database (Wong 2010) |
| | GC content | UCSC |
| regulation | CpG islands | UCSC |
| functional | refseq Exons | NCBI |
| | refseq Genes | NCBI |
| conservation | SV | DGV |
| | selfChain | UCSC |
| | Fungtammasan CommonFragileSites | Fungtammasan(2012) |
| | Syntenic blocks | results 1_1 |
| | CEGA_Amniota | CEGA |
| others | RNA G quadruplex | G4RNA |

**Table S6.** Genomic features description

| | BHR | | BRR | | Other | |
|---|---|---|---|---|---|---|
| **Feature** | **rank** | **coefficient** | **rank** | **coefficient** | **rank** | **coefficient** |
| **RNA G4** | 0 | 0.000187875 | 11 | -2.06418e-6 | 23 | -0.000185811 |
| **Tandem Repeat** | 1 | 0.000185345 | 10 | -1.93254e-6 | 22 | -0.000183412 |
| **SINE** | 2 | 0.000137291 | 23 | -0.000244478 | 2 | 0.000107186 |
| **G4 forming Repeat** | 3 | 0.0000740295 | 7 | 8.01735e-6 | 20 | -0.000082047 |
| **Direct Repeat** | 4 | 0.0000738793 | 22 | -0.000114042 | 7 | 0.0000401623 |
| **Mirror Repeat** | 5 | 0.0000623136 | 20 | -0.000103197 | 6 | 0.0000408831 |
| **Synteny Block** | 6 | 0.0000394735 | 19 | -0.000090527 | 4 | 0.0000510532 |
| **Simple Repeat** | 7 | 0.0000371608 | 15 | -1.71908e-5 | 19 | -0.00001997 |
| **G4** | 8 | 0.0000203475 | 21 | -0.000110452 | 3 | 0.0000901043 |
| **refSeq Exon** | 9 | 0.0000142889 | 17 | -0.000043113 | 10 | 0.0000288239 |
| **Common Fragile Site** | 10 | 5.77519e-6 | 14 | -1.30962e-5 | 14 | 0.000007321 |
| **DGV Structural Variation** | 11 | 1.26402e-6 | 13 | -4.79724e-6 | 16 | 3.53322e-6 |
| **refSeq Gene** | 12 | 7.48958e-7 | 12 | -4.29641e-6 | 15 | 3.54745e-6 |
| **Self Chain** | 13 | -5.35611e-7 | 8 | 2.54847e-6 | 17 | -2.01286e-6 |
| **LINE** | 14 | -8.1989e-6 | 16 | -2.80981e-5 | 8 | 0.000036297 |
| **Segmental Duplication** | 15 | -8.62652e-6 | 9 | -6.23996e-7 | 12 | 9.25052e-6 |
| **LTR** | 16 | -0.000041455 | 5 | 0.0000341031 | 13 | 7.35197e-6 |
| **CpG island** | 17 | -0.000055191 | 6 | 0.0000112632 | 5 | 0.0000439274 |
| **Satellite** | 18 | -0.000109061 | 4 | 0.0000776391 | 9 | 0.0000314215 |
| **ZDNA** | 19 | -0.000112429 | 18 | -0.000075118 | 1 | 0.000187547 |
| **Inverted Repeat** | 20 | -0.000117265 | 3 | 0.0000996355 | 11 | 0.0000176298 |
| **CEGA Amniota** | 21 | -0.000158965 | 2 | 0.000278049 | 21 | -0.000119084 |
| **Low Complx Repeat** | 22 | -0.00042357 | 1 | 0.00044245 | 18 | -1.88794e-5 |
| **a-Phased Repeat** | 23 | -0.00111544 | 0 | 0.000488061 | 0 | 0.000627379 |

**Table S7.** Logistic Regression Coefficient

| | Coverage in percentage | | |
|---|---|---|---|
| **Genomic feature** | **genome** | **EBHR** | **EBRR** |
| DGV SV | 79.45 | 87.00 | 14.55 |
| refseq Genes | 20.82 | 23.00 | 1.36 |
| self Chain | 20.14 | 23.10 | 33.13 |
| Syntenic blocks | 52.52 | 56.07 | 0.72 |
| Common Fragile Sites | 15.13 | 15.12 | 0.13 |
| CEGA Amniota | 2.58 | 2.38 | 0.08 |
| inverted repeats | 4.13 | 4.49 | 0.67 |
| LINE | 10.04 | 9.30 | 1.17 |
| SD | 3.73 | 2.73 | 4.65 |
| LTR | 3.11 | 3.04 | 0.93 |
| mirror repeats | 2.45 | 2.81 | 0.46 |
| direct repeats | 1.33 | 1.64 | 0.35 |
| short tandem repeats | 1.39 | 1.69 | 0.25 |
| refseq Exons | 1.17 | 1.37 | 0.09 |
| SINE | 0.93 | 1.35 | 0.06 |
| CpG island | 0.68 | 0.81 | 0.08 |
| Satellite repeats | 0.13 | 0.02 | 1.21 |
| a-phased repeats | 0.33 | 0.32 | 0.08 |
| g-quadruplex-forming repeats | 0.16 | 0.20 | 0.03 |
| Z-DNA | 0.20 | 0.23 | 0.03 |
| G quadruplex | 0.32 | 0.42 | 0.04 |
| RNA G quadruplex | 0.00 | 0.01 | 0.00 |
| Low complexity repeats | 0.02 | 0.03 | 0.02 |
| Simple repeat | 0.14 | 0.22 | 0.07 |

**Table S8.** The coverage of different genomic regions by selected features
